# BiMultiNetPlot: An R package for visualizing ecological bipartite multilayer networks

**DOI:** 10.1101/2024.09.20.613870

**Authors:** Hai-Dong Li

**Author notes:** **Correspondence** Hai-Dong Li.

## Abstract

1. With the increasing study of ecological multilayer bipartite networks, the visualization of these networks has become more important. However, tools for visualizing multilayer networks are still lacking.
2. I present BiMultiNetPlot, an R package designed for visualizing ecological bipartite multilayer networks.
3. I demonstrate how to use BiMultiNetPlot through a series of examples that represent the most common types of ecological multilayer networks.
4. BiMultiNetPlot is an open-source, flexible package within the ggplot2 environment, helping ecologists better understand the multilayer nature of ecological networks.

## 1 INTRODUCTION

Species and individuals within ecological systems interact with each other, and these interactions form ecological networks (Bascompte & Jordano 2007). Biodiversity maintenance and ecosystem functioning depend upon these interactions (Bascompte & Jordano 2013; Schleuning, Fruend & Garcia 2015). The graphical presentation of networks, depicting who interacts with whom, is an essential component of ecological network studies, as it is inherently engaging and intuitively interpretable and plays a crucial role in scientific dissemination (Pocock *et al*. 2016). So far, there are many R packages available for visualizing ecological networks, such as bipartite (Dormann, Gruber & Fründ 2008), bipartiteD3 (Terry 2024) igraph (Csardi & Nepusz 2006), and so on. These R packages have definitively promoted the area of ecological network studies. However, such tools are generally designed for visualizing monolayer networks and are not very well-suited for representing the ecological multilayer networks that have been proposed recently (Pilosof *et al*. 2017; Hutchinson *et al*. 2018). Therefore, there is a need to develop tools for ecological multilayer network visualization.

Multilayer networks, providing a more realistic representation of complex ecological systems, have multiple “layers” and interlayer links connecting nodes from different layers, and the network in each layer is a monolayer network (Pilosof *et al*. 2017). Layers typically represent space (Timoteo *et al*. 2018), time (Costa *et al*. 2020), and interaction types (Mello *et al*. 2019). In spatial and temporal ecological multilayer networks, layers are space and time points, respectively. They consist of multiple networks of a focal species interaction type (e.g., plant-pollinator interactions) connected via inter-layer links that describe an additional ecological process, such as species movement between habitats or population dynamics through time (Fig. 1A and B). In multilayer networks that are composed of different types of species interactions (e.g., pollination, seed dispersal), each ecological interaction occurs in a different layer, and interlayer links connect shared species to their counterparts in other layers (Fig. 1C). Such multilayer networks are typically ‘diagonally coupled’, thus called diagonal coupling multilayer networks (and also called multiplex networks). Biologically, interlayer links in multiplex networks may represent the influence of one interaction type on the other. With the increasing study on ecological multilayer networks (Timoteo *et al*. 2018; Mello *et al*. 2019; Costa *et al*. 2020; Hervias-Parejo *et al*. 2020; Hervías-Parejo *et al*. 2023; Vitali *et al*. 2023), the visualization of ecological bipartite multilayer networks become more important.

**Figure 1.**
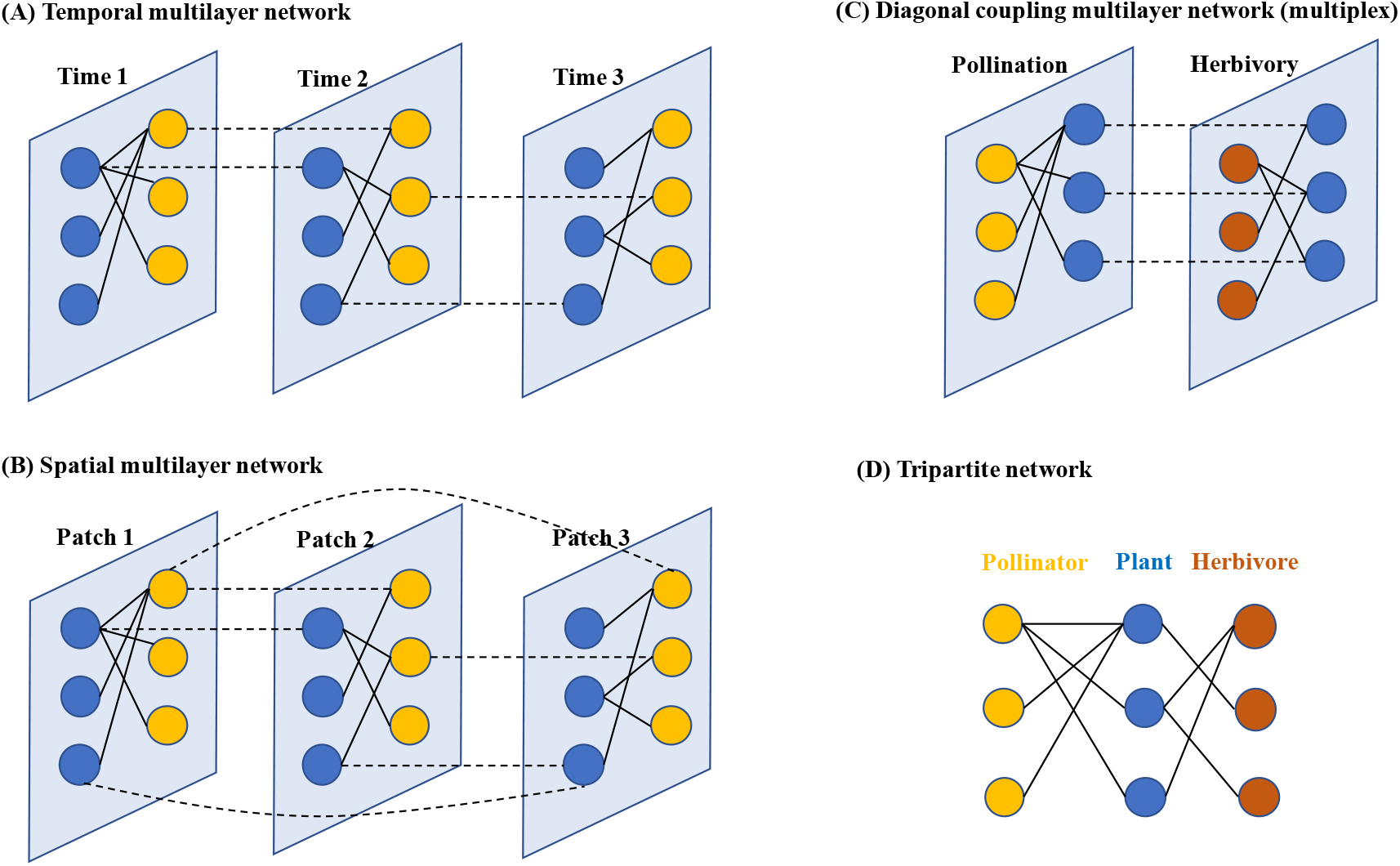
The examples of multilayer bipartite networks. (A) A temporal multilayer network that contains three layers and two levels of species (i.e., blue and yellow points). (B) A spatial multilayer network that contains three layers and two levels of species (i.e., blue and yellow points). (C) A diagonal coupling multilayer network (or called multiplex network) that contains two interaction types (pollination and herbivory) and three levels of species (i.e., blue, brown, and yellow points). In A, B, and C, solid and dashed lines indicate intra-layer links and interlayer links, respectively. Each layer is a bipartite network. (D) A tripartite network that contains three levels of species and it is a diagonal coupling multilayer network that does not have interlayer links.

However, the tools for visualizing multilayer networks are still lacking (but see Hammoud 2018).

To address this knowledge gap, I developed ‘BiMultiNetPlot’, an R package for visualizing bipartite multilayer networks. The BiMultiNetPlot package includes R functions to plot multilayer bipartite networks within the ggplot2 environment. Using the foundational structure of ggplot2, the tidyverse, and related R packages (Wickham 2016; Wickham *et al*. 2019), plotting functions generate ggplot objects with default aesthetics, which can be customized as needed. The package contains functions for visualizing various types of bipartite multilayer networks, including temporal multilayer networks, spatial multilayer networks, diagonal coupling multilayer networks, and tripartite networks. I first provide instructions for installing the package and details on its implementation. Second, I demonstrate the functionality of BiMultiNetPlot through usage examples.

## 2 INSTALLATION

The **BiMultiNetPlot** package can be installed from GitHub via the remotes R package (Csárdi et al., 2023):

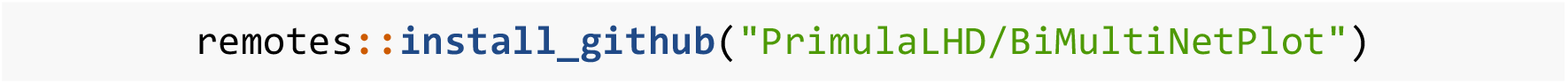

and/or installed from GitHub via the devtools R package (Wickham, Hester, et al., 2022):

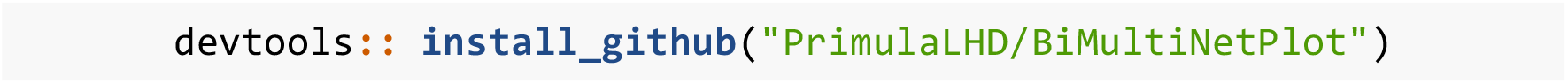

Following installation, **BiMultiNetPlot** can be loaded via the library function in R:

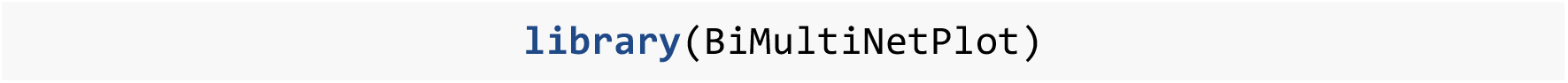

### 2.1 Dependencies

The BiMultiNetPlot package depends on R (> 4.0) (R Core Team, 2024) and imports functions from ggplot2 (Wickham 2016) and tidyverse (Wickham *et al*. 2019). The package was developed with the support of the R packages devtools (Wickham *et al*. 2022), testthat (Wickham 2011), and roxygen2 (Wickham, Danenberg & Csárdi 2022).

## 3 IMPLEMENTATION AND APPLICATIONS

Table 1 provides a summary of the functions currently available in BiMultiNetPlot. Detailed description is provided by usage examples to demonstrate the functionality of the package. Here I briefly describe the functionality for the five main tasks performed by BiMultiNetPlot (Table 1). I also provide some examples of functions to show the usage of BiMultiNetPlot (Figure 2-6).

**Table 1.** Summary table of the functions currently available in the ‘BiMultiNetPlot’ R package.

| Function | Description |
| --- | --- |
| `plot_temporal_multilayer()` | plots out a temporal multilayer bipartite network |
| `plot_spatial_multilayer()` | plots out a spatial multilayer bipartite network |
| `plot_diagonal_multilayer()` | plots out a diagonal coupling multilayer bipartite network |
| `plot_multiplex` | plots out a general multiplex network |
| `plot_tripartite()` | plots out a tripartite network |
| `plot_monolayer()` | plots out a bipartite network |

**Figure 2.**
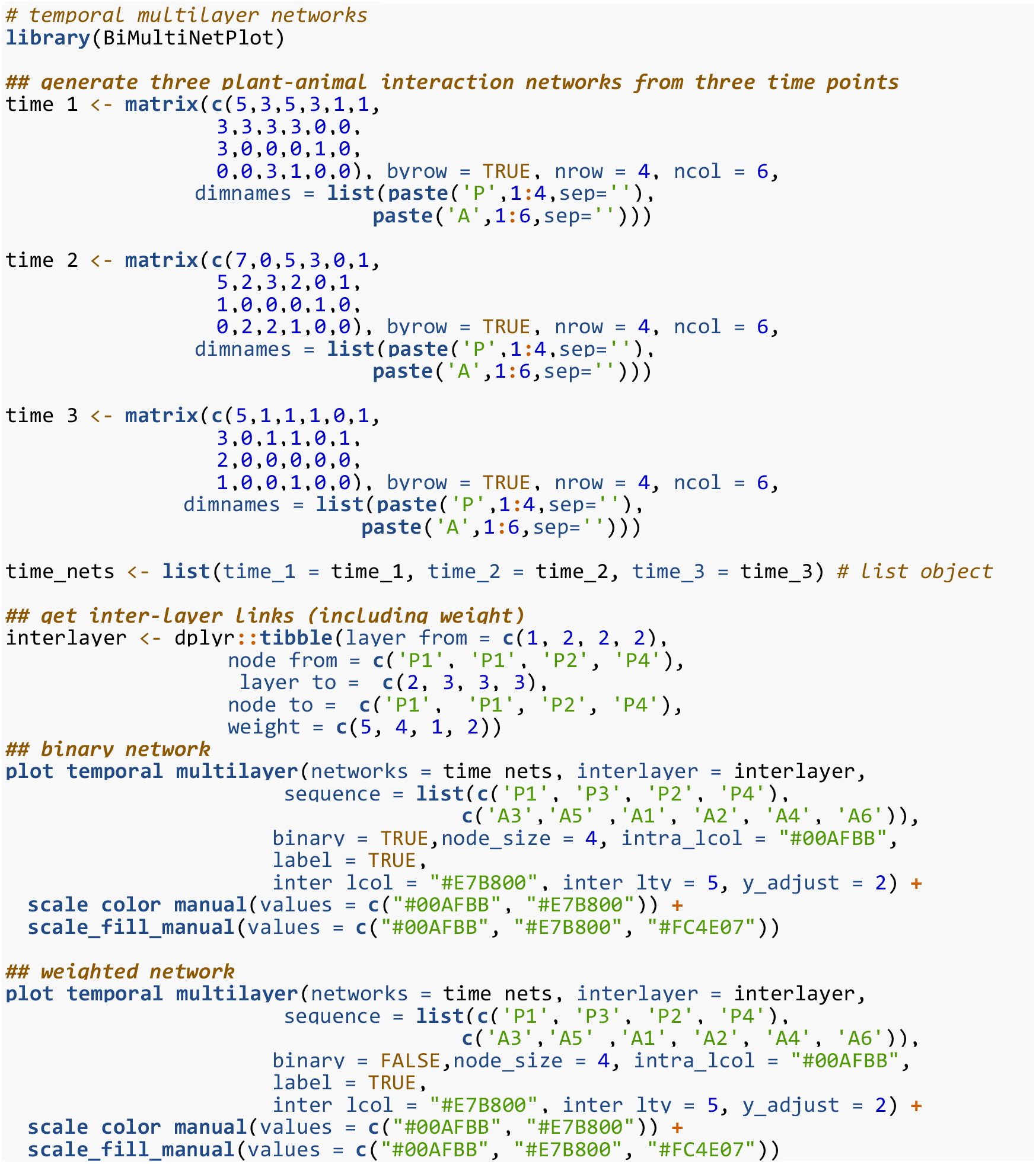
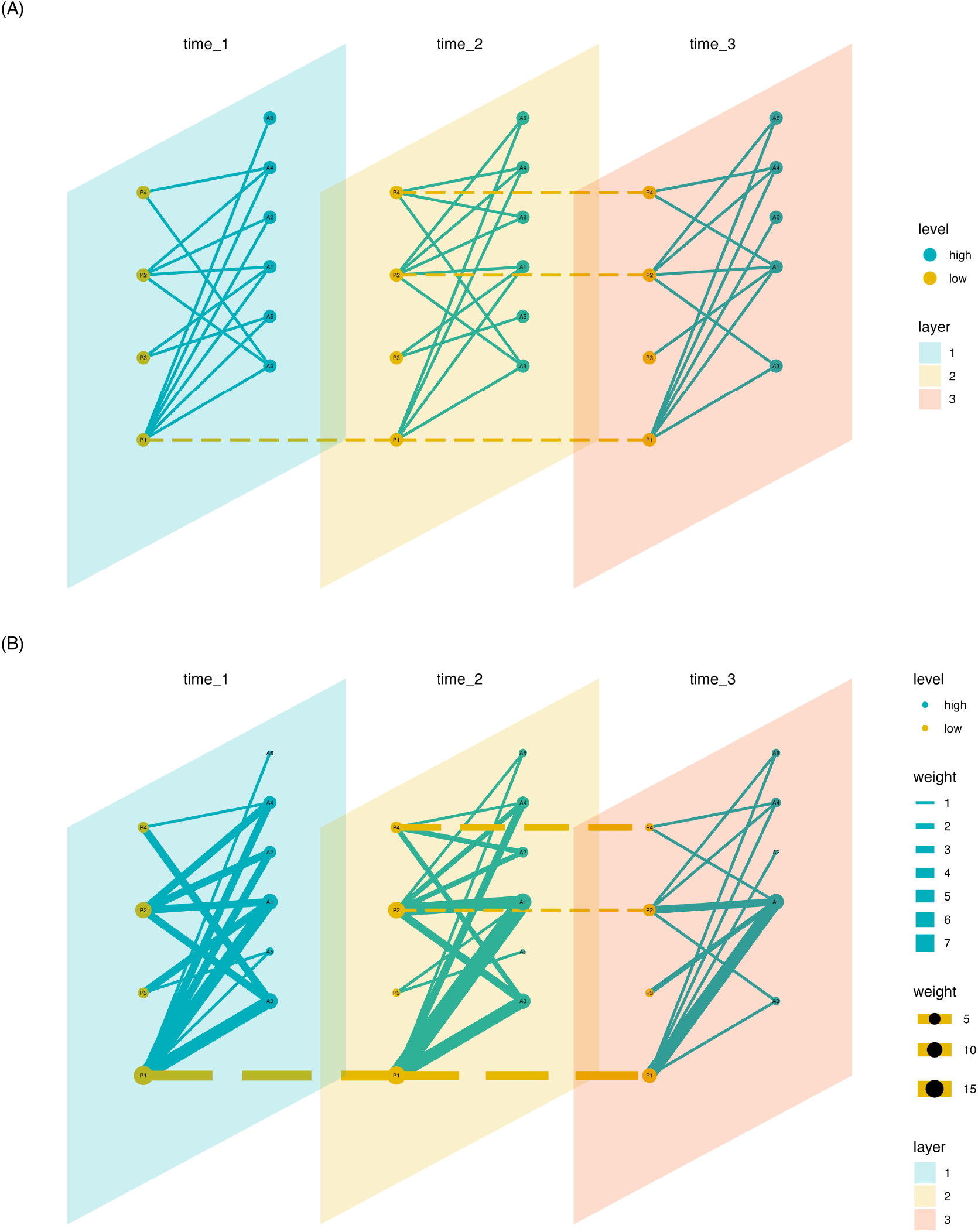
Examples of the temporal multilayer network of plant-animal interactions. (A) binary networks, and (B) weighted networks. Green lines and brown lines indicate intra-layer links and inter-layer links, respectively. Node size in the weighted network indicates the interaction strength of the intra-layer, and the width of lines indicates link strength.

### 1. Temporal multilayer network

The function *plot_temporal_multilayer* visualizes temporal multilayer bipartite networks using the ggplot2 style. Users should provide a list object of species interaction matrix and an interlayer link data frame (describing which species connected from one layer to another layer). In each interaction matrix, low trophic species are rows and high trophic species are columns. Users can specify each parameter to customize plots’ aesthetics. BiMultiNetPlot also allows users to provide an extended data frame to modify the node’s color attribute. For example, this is useful in the case that users want to show species module affiliation.

To demonstrate the functionality of plotting a temporal multilayer network, I used simulated data and plotted a binary and a weighted multilayer network by setting the parameter “binary”, respectively (Figure 2):

### 2. Spatial multilayer network

The function *plot_spatial_multilayer* visualizes spatial multilayer bipartite networks using the ggplot2 style (Figure 3). Similar to temporal multilayer networks, users should provide a list object of species interaction matrix, an inter-layer link data frame, and also could provide an extended data frame. Different from temporal multilayer networks, the layers are not time-ordered, but the layers’ order is not important for spatial multilayer networks. The interlayer links are unidirectional in temporal multilayer networks, but they are generally non-directional in spatial multilayer networks. Simulated data and spatial multilayer bipartite network were shown with examples in Figure 3.

**Figure 3.**
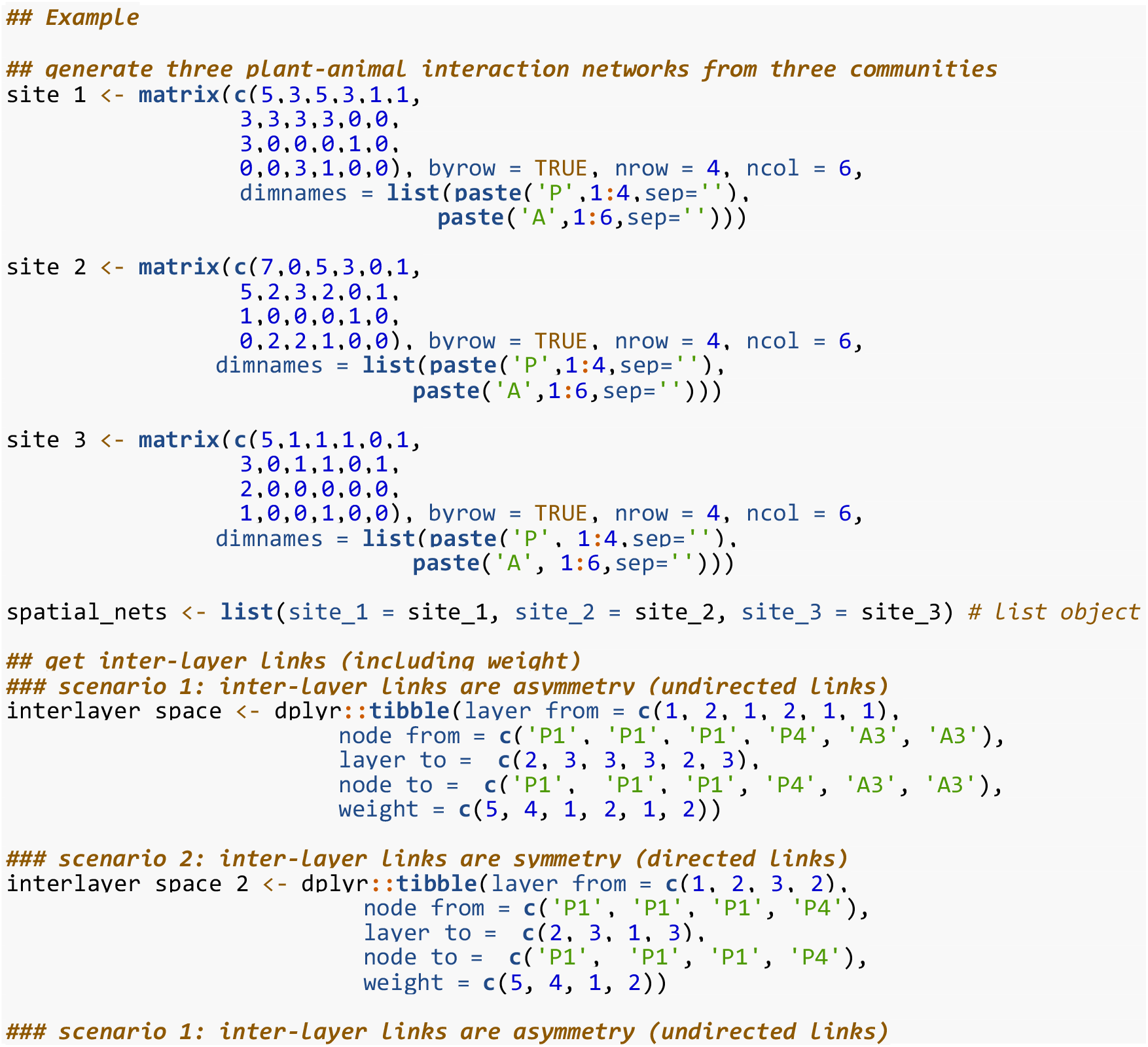
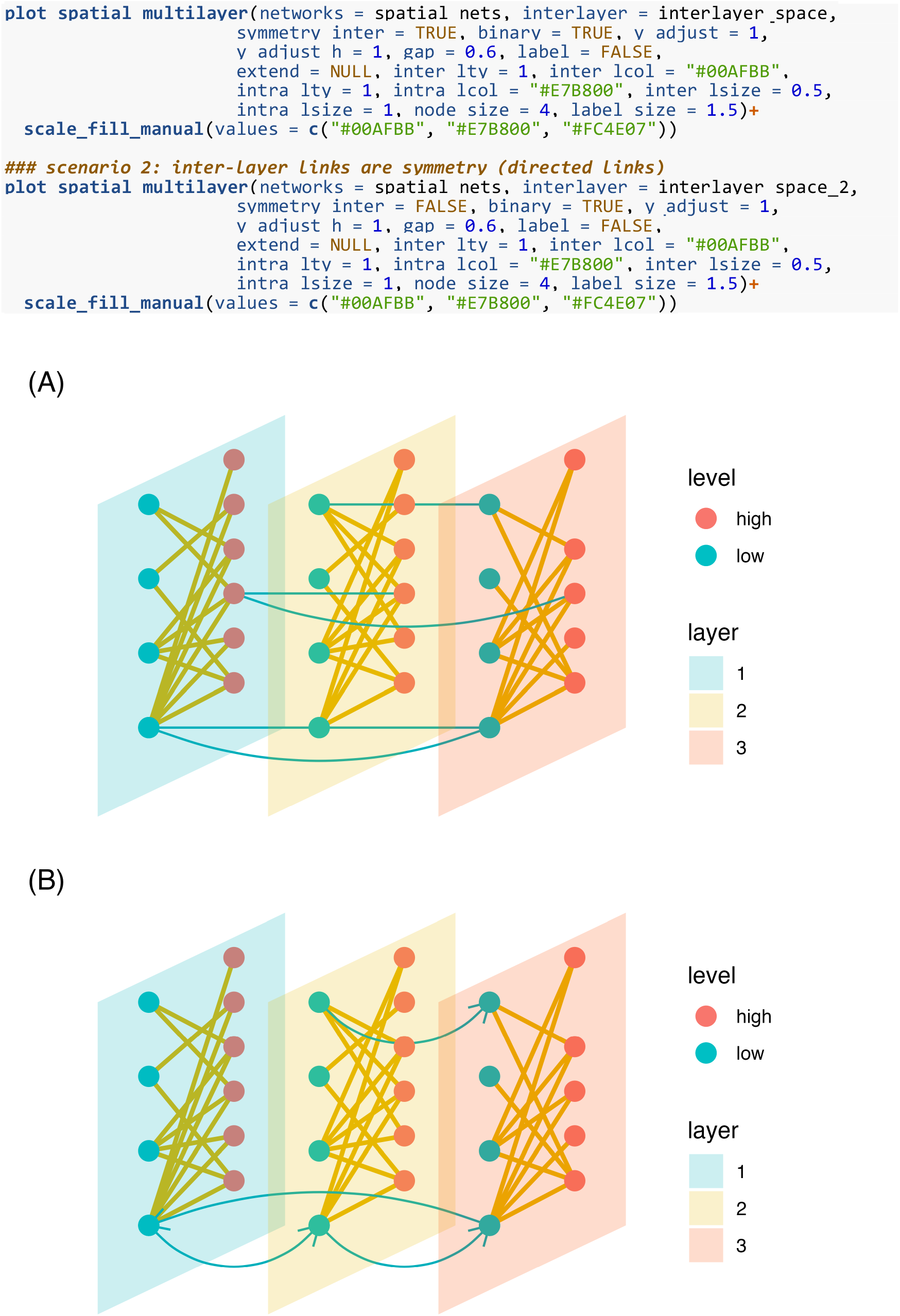
Examples of the spatial multilayer network of plant-animal interactions. (A) unidirectional binary networks, and (B) directional binary networks. Green lines and brown lines indicate inter-layer links and intra-layer links, respectively. Lines with arrows in the directional network indicate the direction of the inter-layer links.

### 3. Diagonal coupling multilayer network

The function *plot_diagonal_multilayer* visualizes diagonal coupling multilayer bipartite networks using the ggplot2 style. Two interaction matrices and an inter-layer links data frame should be provided. Other aesthetics and options are similar to functions *plot_temporal_multilayer* and *plot_spatial_multilayer*. Simulated data and diagonal coupling multilayer bipartite network were shown with example in Figure 4.

**Figure 4.**
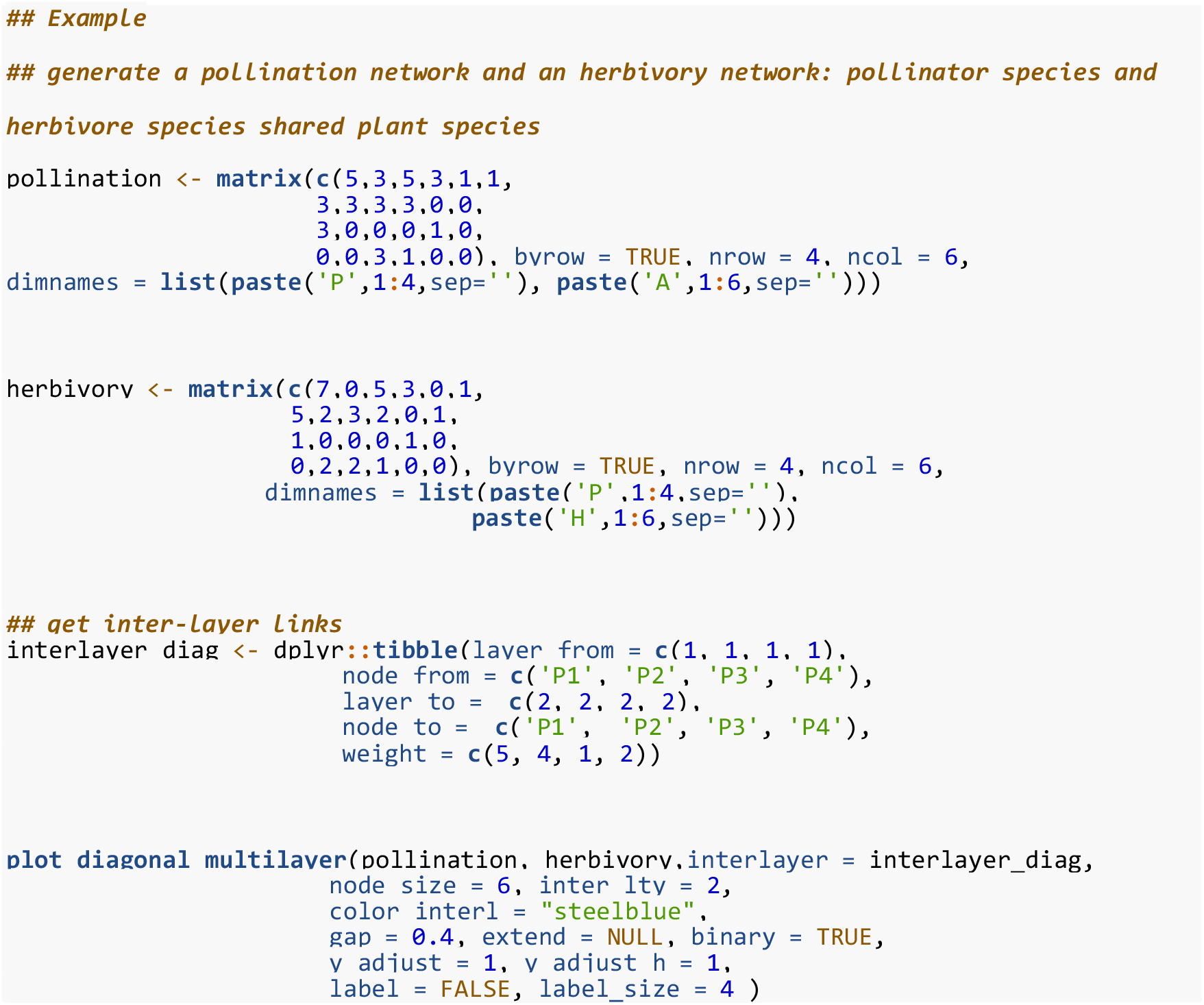
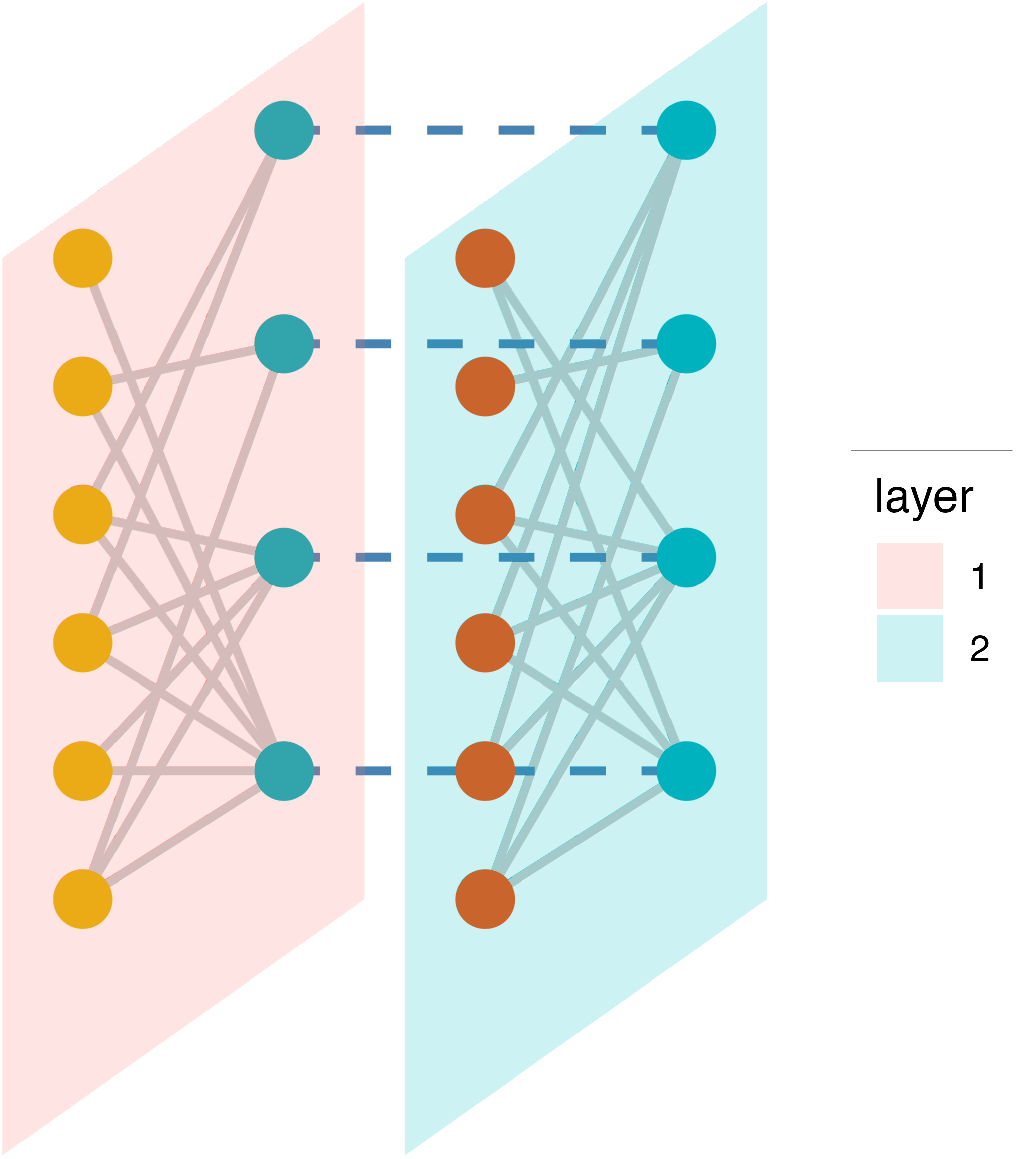
An example of the diagonal coupling multilayer network. The network on the left is the pollination network, and the right is the herbivory network.

### 4. Tripartite network

The function *plot_tripartite* visualizes tripartite networks using the ggplot2 style. A tripartite network is a diagonal coupling multilayer network that does not have interlayer links. Users should provide two interaction matrices and could provide an extended data frame additionally. The two interaction matrices represent two interaction types, and the common species are rows in the two matrices. Simulated data and tripartite network were shown with example in Figure 5.

**Figure 5.**
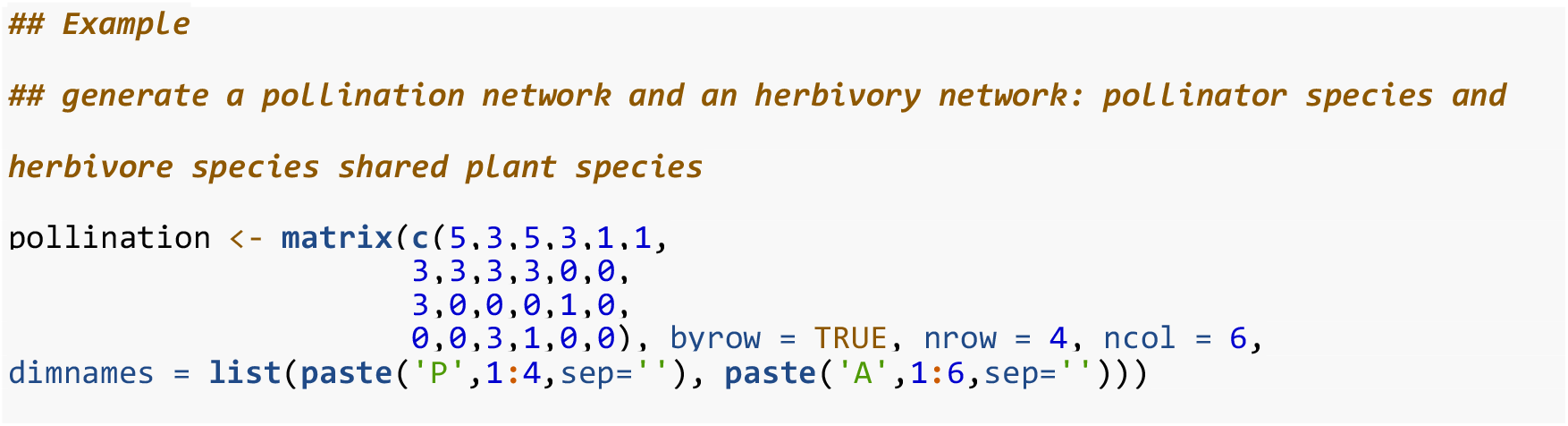
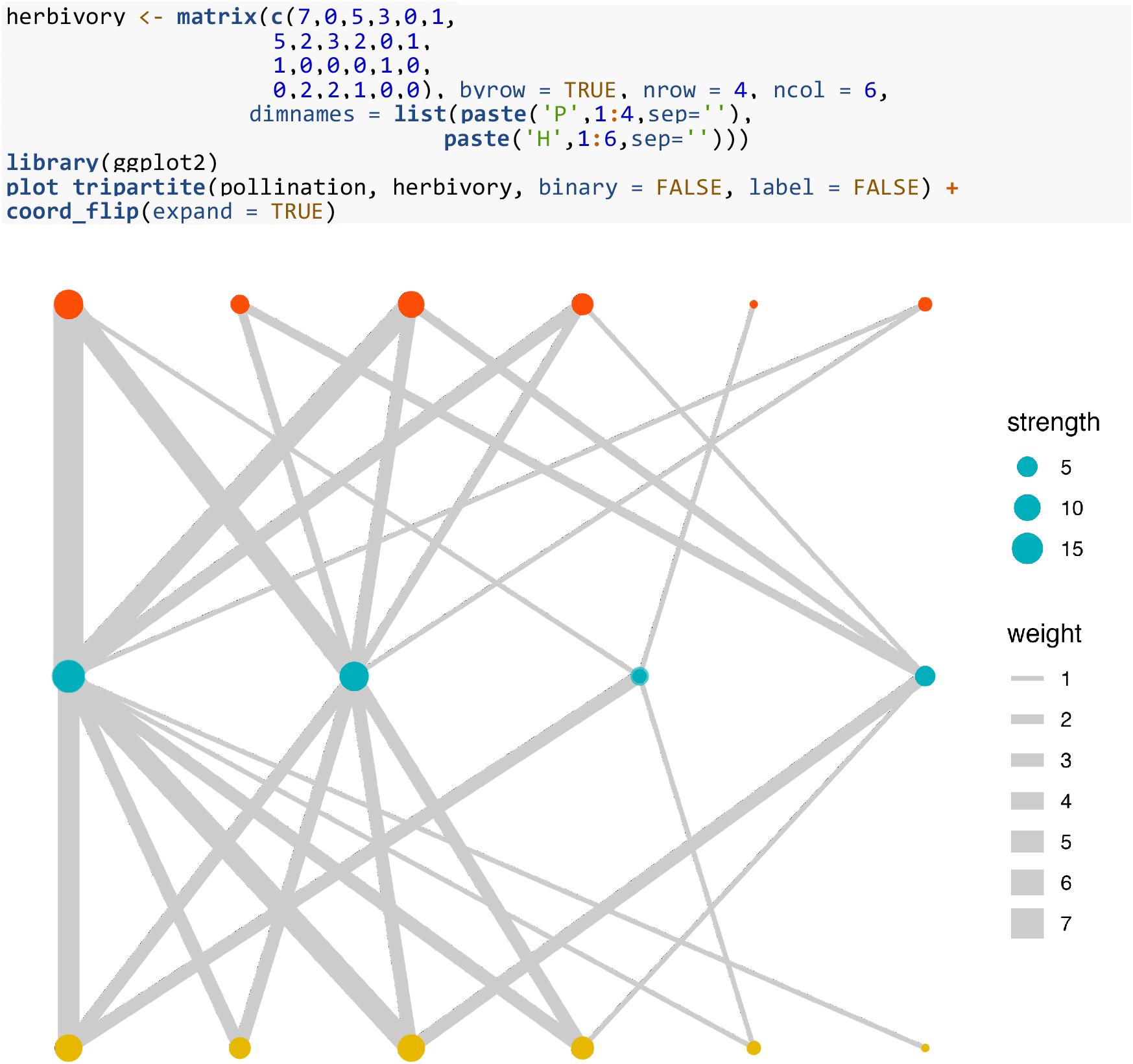
An example of the tripartite network. The nodes at the bottom are pollinators, the nodes at the middle are plants, and the nodes at the top are herbivores.

### 5. Monolayer bipartite network

The function *plot_monolayer* visualizes monolayer bipartite networks within the ggplot2 style. To achieve completeness, I also provide this function to plot out a bipartite graph. The input data should be an interaction matrix that low trophic species are rows and high trophic species are columns. Tripartite network was shown with example in Figure 6.

**Figure 6.**
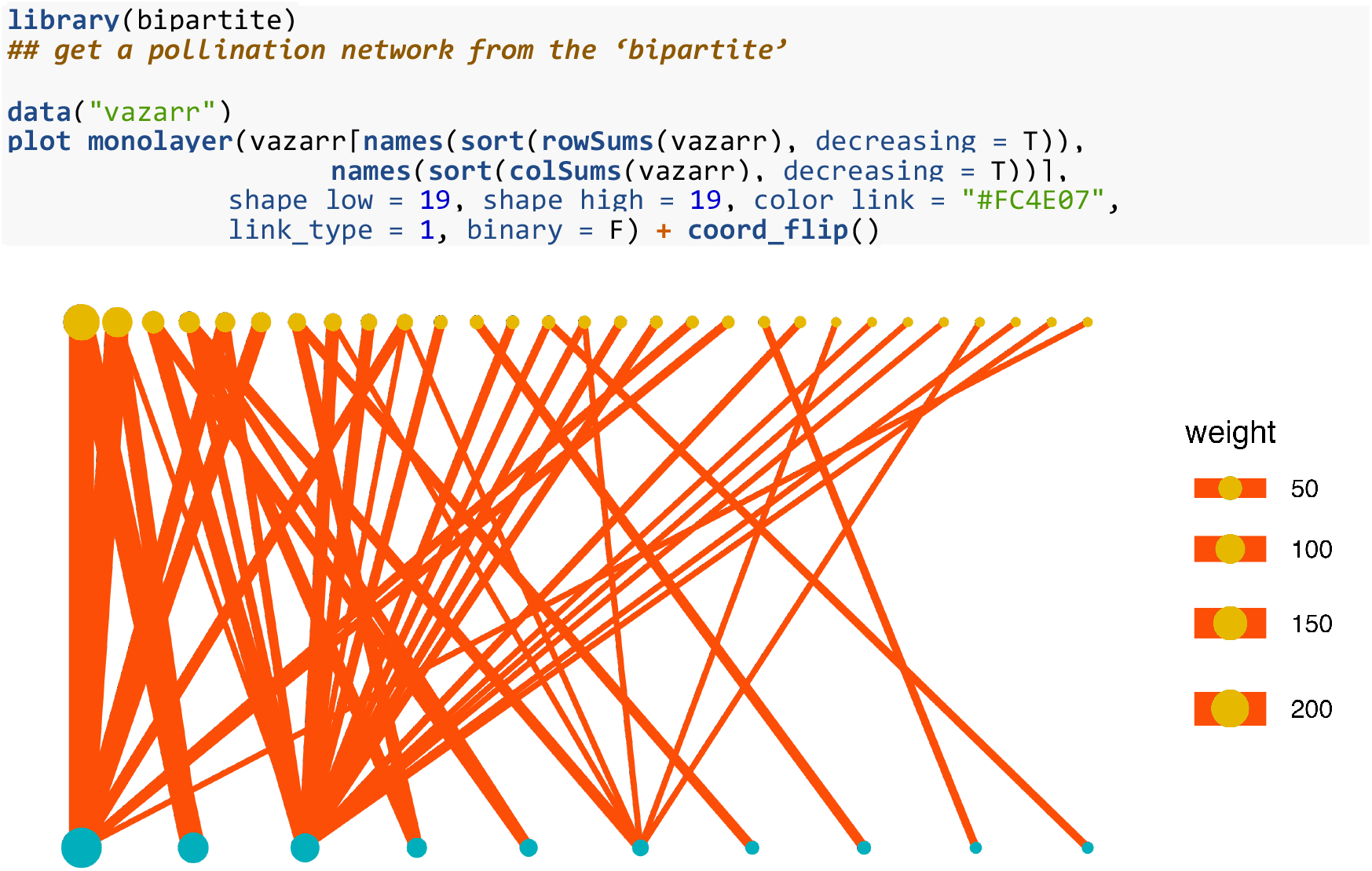
An example of the monolayer pollination network. Data was from the “bipartite” package (Dormman et al. 2008). The nodes at the bottom are plants, and the nodes at the top are animals.

## 4 CONCLUSIONS

The BiMultiNetPlot R package fits multilayer bipartite networks. The package includes functions for returning temporal multilayer networks, spatial multilayer networks, tripartite networks, and monolayer networks. I expect that BiMultiNetPlot will serve as a user-friendly tool for ecologists who interested in the multilayer nature of ecological networks.

## ACKNOWLEDGEMENTS

This work was supported by the National Natural Science Foundation of China (32101284).

## CONFLICTS OF INTEREST

The authors declare no conflicts of interest.

## AUTHORS’ CONTRIBUTIONS

H.D.L designed the R package and drafted the manuscript.

## DATA AVAILABILITY STATEMENT

BiMultiNetPlot package is available on GitHub (https://github.com/PrimulaLHD/BiMultiNetPlot). All examples are freely available and are bundled with the R package.

## Notes

### Competing Interest Statement

The authors have declared no competing interest.

